# Inferring extrinsic noise from single-cell gene expression data using Approximate Bayesian Computation

**DOI:** 10.1101/030155

**Authors:** Oleg Lenive, Paul DW Kirk, Michael PH Stumpf

## Abstract

**Background:** Gene expression is known to be an intrinsically stochastic process which can involve single-digit numbers of mRNA molecules in a cell at any given time. The modelling of such processes calls for the use of exact stochastic simulation methods, most notably the Gillespie algorithm. However, this stochasticity, also termed “intrinsic noise”, does not account for all the variability between genetically identical cells growing in a homogeneous environment. Despite substantial experimental efforts, determining appropriate model parameters continues to be a challenge. Methods based on approximate Bayesian computation can be used to obtain posterior parameter distributions given the observed data. However, such inference procedures require large numbers of simulations of the model and exact stochastic simulation is computationally costly. In this work we focus on the specific case of trying to infer model parameters describing reaction rates and extrinsic noise on the basis of measurements of molecule numbers in individual cells at a given time point.

**Results:** To make the problem computationally tractable we develop an exact, model-specific, stochastic simulation algorithm for the commonly used two-state model of gene expression. This algorithm relies on certain assumptions and favourable properties of the model to forgo the simulation of the whole temporal trajectory of protein numbers in the system, instead returning only the number of protein and mRNA molecules present in the system at a specified time point. The computational gain is proportional to the number of protein molecules created in the system and becomes significant for systems involving hundreds or thousands of protein molecules. We employ this algorithm, approximate Bayesian computation, and published gene expression data for *Escherichia coli* to simultaneously infer the model’s rate parameters and parameters describing extrinsic noise for 86 genes.

## Background

Experiments have demonstrated the presence of considerable cell-to-cell variability in mRNA and protein numbers^1–5^ and slow fluctuations on timescales similar to the cell cycle.^6,7^ Broadly speaking, there are two plausible causes of such variability. One is the inherent stochasticity of biochemical processes which are dependent on small numbers of molecules. The other relates to differences in numbers of protein, mRNA, metabolites and other molecules available for each reaction or process within a cell, as well as any heterogeneity in the physical environment of the cell population. These sources of variability have been dubbed as “intrinsic noise” and “extrinsic noise”, respectively.

One of the earliest investigations into the relationship between intrinsic and extrinsic noise employed two copies of a protein with different fluorescent tags, expressed from identical promoters equidistant from the replication origin in *E. coli.*^8^ By quantifying fluorescence for a range of expression levels and genetic backgrounds the authors concluded that intrinsic noise decreases monotonically as transcription rate increases while extrinsic noise attains a maximum at intermediate expression levels. Other studies have considered extrinsic noise in the context of a range of cellular processes including the induction of apoptosis;^9^ the distribution of mitochondria within cells;^10^ and progression through the cell cycle.^11^ From a computational perspective, extrinsic variability has been modelled by linking the perturbation of model parameters to perturbation of the model output using the Unscented Transform.^12^

Taniguchi *et al*^7^ carried out a high-throughput quantitative survey of gene expression in *E. coli.* By analysing images from fluorescent microscopy they obtained discrete counts of protein and mRNA molecules in individual *E. coli* cells. They provided both the measurements of average numbers of protein and mRNA molecules in a given cell, as well as measurements of cell-to-cell variability of molecule numbers. The depth and scale of their study revealed the influence of extrinsic noise on gene expression levels. The authors demonstrated that the measured protein number distributions can be described by Gamma distributions, the parameters of which can be related to the transcription rate and protein burst size.^13^ To quantify extrinsic noise they consider the relationship between the means and the Fano factors of the observed protein distributions. They also illustrate how extrinsic noise in protein numbers may be attributed to fluctuations occurring on a timescale much longer than the cell cycle.

Here we aim to describe extrinsic noise at a more detailed, mechanistic, level using a stochastic model of gene expression. Such a description calls for quantitative inference of the model’s parameters. We achieve this by relying on the data made available by Taniguchi *et al* and employing approximate Bayesian computation (ABC).^14,15^ One difficulty that arises when trying to investigate the extent and effect of extrinsic noise is that it is difficult to separate it from intrinsic noise. To overcome this confounding effect, the parameters of our model come in two varieties. Firstly, reaction rate parameters describe the probability of events occurring per unit of time. These correspond to the reaction rate parameters of a typical stochastic model which accounts for intrinsic noise. Secondly, noise parameters describe the variability in reaction rate parameters caused by the existence of extrinsic noise. This approach allows us to simultaneously infer the rate parameters and the magnitude of extrinsic noise.

Stochastic simulation and ABC inference methods are both computationally costly endeavours. In this particular case, the experimental data corresponds to snapshots of the system at a single time point. Thus, a complete temporal trajectory of the system is not necessary to carry out comparisons with the data. This allows us to make the problem computationally tractable. To this end, we develop a model-specific simulation method which takes advantage of the Poissonian relationship between the number of surviving protein molecules produced from a given mRNA molecule and its lifetime, under certain assumptions.

## Results and discussion

### Posterior distributions of parameters

We begin our analysis by examining the posterior distributions of parameters obtained for each gene using the ABC-SMC inference procedure.^14^ The simulated summary statistics converged to within the desired threshold of the experimental measurements for 86 out of 87 genes. The inferred posterior for the one remaining gene converged relatively slowly and we chose to terminate the process after 30 days of CPU time.

Figure 1 shows a contour plot of the distribution of summary statistics and the mRNA degradation rate, obtained from particles in the final ABC-SMC population for a typical gene (*dnaK*). We begin with a discussion of features of the posterior parameter distributions, that are common to most genes. Next, we examine the relationships between model parameters and summary statistics of the model outputs. Lastly, we carry out a sensitivity/robustness analysis on the inferred posteriors to assess the importance of each parameter in setting the overall levels of extrinsic noise.

**Figure 1.**
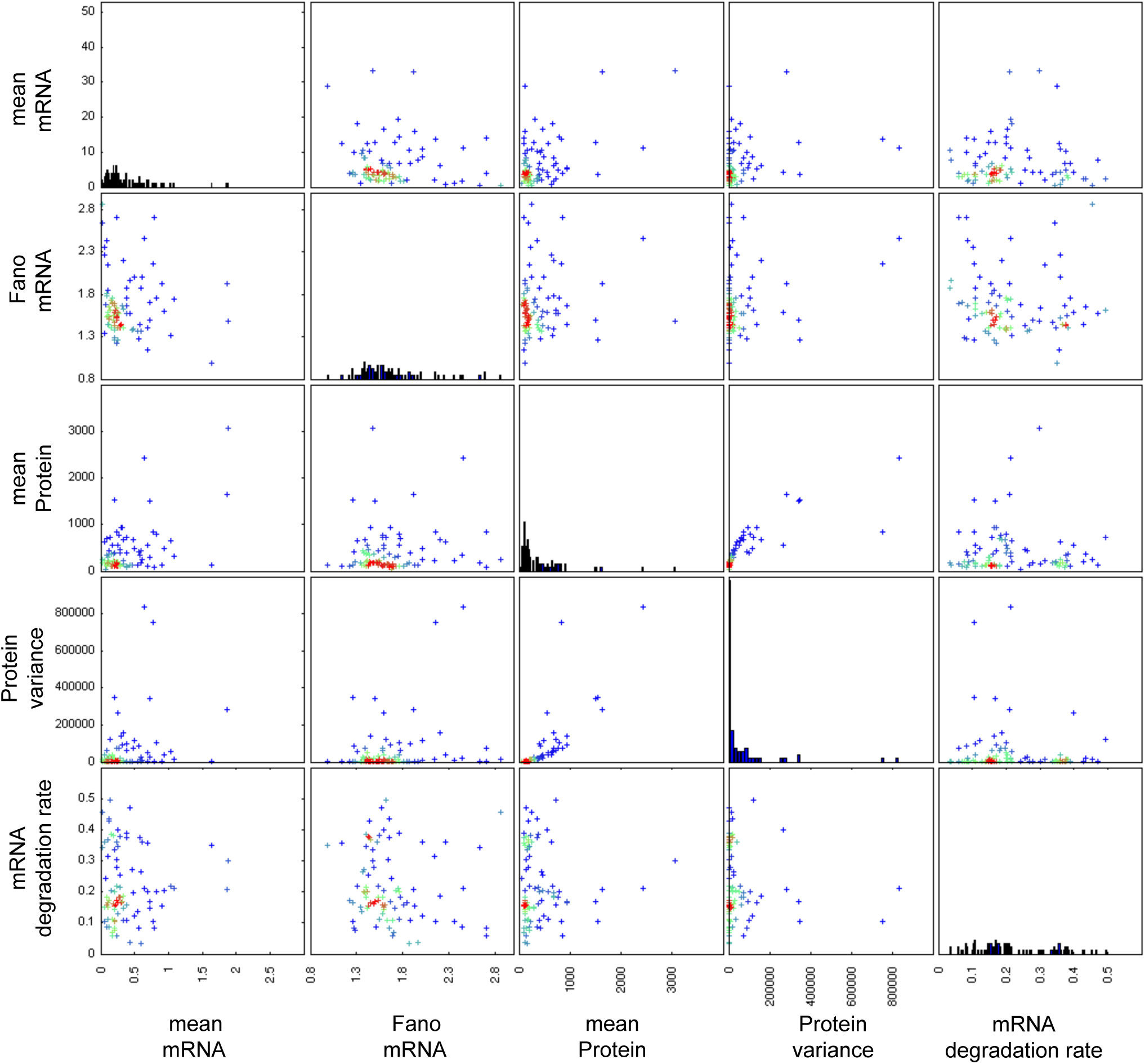
Posterior distribution of summary statistics for the gene *dnaK.* Contour plots indicating the density of points with the corresponding summary statistic for each particle in the final population.

In the two-state model, the switching of the promoter between active and inactive states is described by a telegraph process that can be parametrised either in terms of the switching reaction rates (*k*_on_ and *k*_off_) or in terms of the on/off bias (*k_r_*) and frequency of switching events (*k_f_*). The simulation algorithm takes parameters in the form of *k*_on_ and *k*_off_. However, the effects of *k_r_* and *k_f_* on the observed mRNA distribution may be interpreted more directly and intuitively.

For the majority of genes the *k*_0_ and *k_r_* parameters are relatively small. This appears to be a prerequisite for a high Fano factor of the mRNA distribution and the mean marginal inferred values of these parameters are negatively correlated with Fano factors across all 86 genes as discussed below. A low switching rate combined with a low basal expression rate ensures that there are two distinct mRNA expression levels. This in turn produces a larger variance in measured mRNA counts and results in Fano factor values well above one. Conversely, genes for which mRNA production appears to be more Poissonian were inferred to have basal mRNA production rates close to one, i.e. similar to the active mRNA production rates. In other words, these genes appear to be constitutively active. Here again, we point out that the two-state promoter model provides a convenient abstraction and a hypothesis for explaining the super-Poissonian variance in mRNA copy number.^5,16^ However, based on these observations it is difficult to determine whether a model with more states or some other more elaborate regulatory model, would not be more appropriate. Our attempts at carrying out the inference procedure with a one-state model indicate that extrinsic noise alone does not explain the observed mRNA distributions without also producing unacceptably high variability in protein numbers.

Our initial inference attempts used only the summary statistics from the data. We observed that the production and degradation rate parameters for mRNA (*k*_1_ and *d*_1_) and protein (*k*_2_ and *d*_2_) tended to be positively correlated in the posterior parameter distributions of many genes. This is due to limited identifiability of model parameters since different combinations of rates may produce similar steady state expression levels. This problem was partly alleviated by directly incorporating information about mRNA degradation in the inference procedure in order to constrain the range of acceptable values for *d*_1_ for each gene (Figure 2). Our approach does provide an indication of the possible range of extrinsic noise values that can account for the observed variability in mRNA and protein numbers.

**Figure 2.**
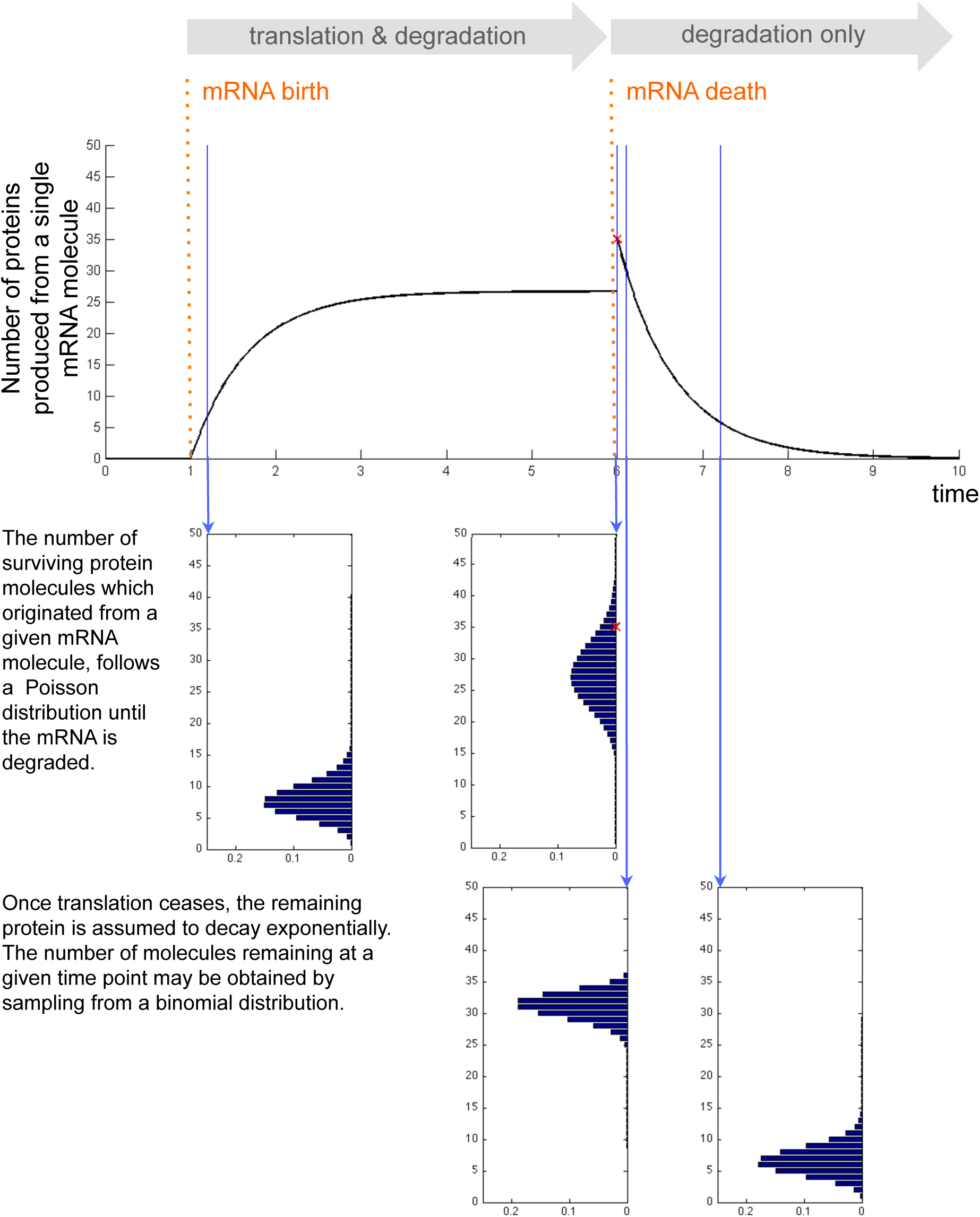
Posterior distribution of model parameters for the gene *dnaK.* Contour plots indicating the density of points with the corresponding parameter values for each particle in the final population.

Although the posterior summary statistics (and mRNA degradation rate) are reasonably well constrained and distinct for each gene, the distributions of model parameters can still be relatively broad (Figure 2). There are a number of reasons for this. Firstly, changes in parameters associated with active transcription and translation, as well as degradation rates, are more easily inferred than parameters describing switching between promoter states, basal transcription or extrinsic noise. In particular, when the production and degradation rates for the same species are subjected to different extrinsic noise parameters, the inference procedure struggles to resolve between the different source of extrinsic noise. This explains the correlation between the means of inferred extrinsic noise parameters (Figure 3). Such correlations between extrinsic noise parameters are not observed in the posterior of each gene or when taking the single particle with the highest weight from the final population of each gene as in Figure 4.

**Figure 3.**
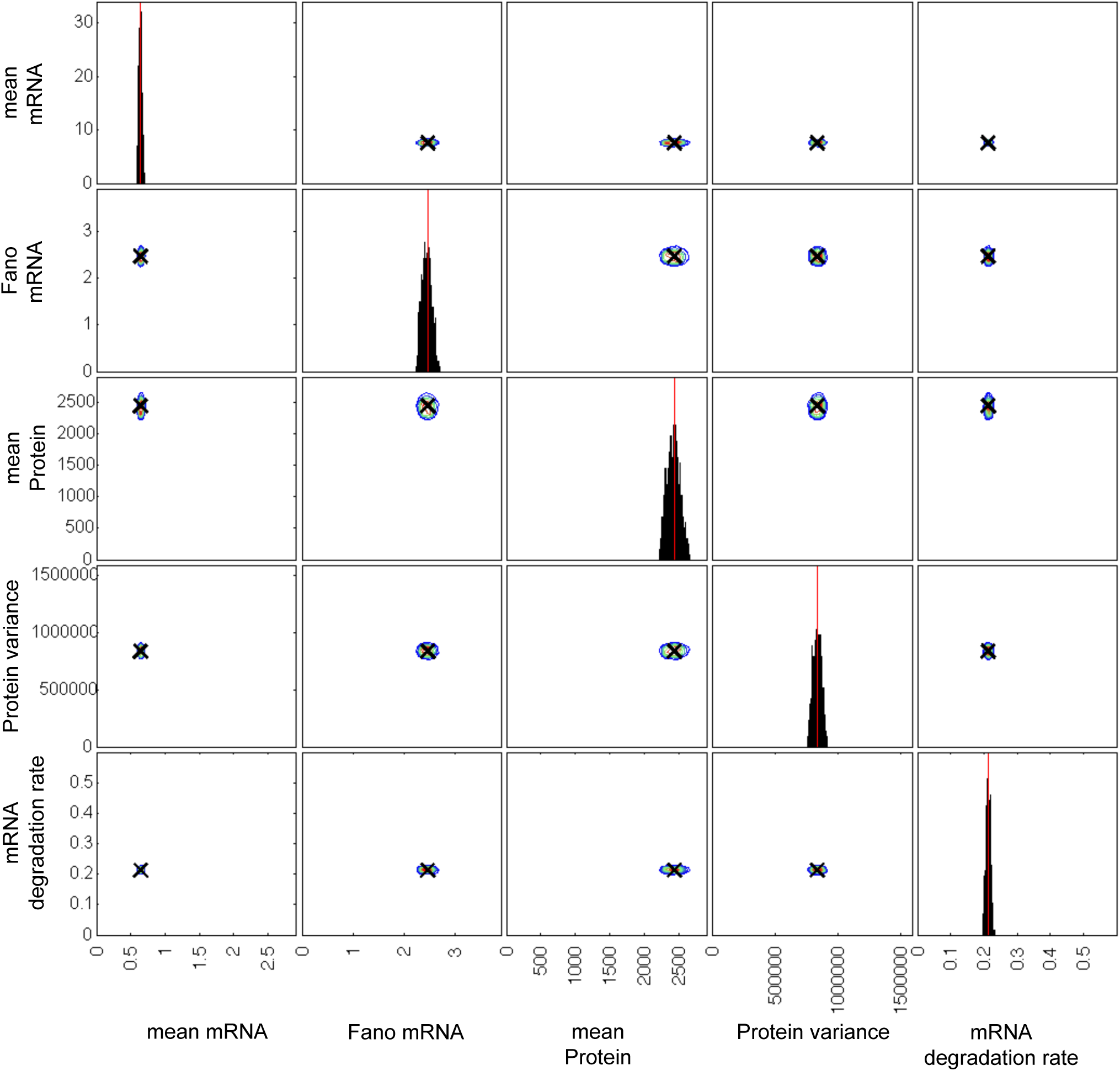
Relationships between means of the marginal parameter posteriors. Scatter plots of the means of the marginal distributions of parameter posteriors are shown for all pairs of parameters. Each point corresponds to a gene. Warmer hues are used to indicate a higher density of data points.

**Figure 4.**
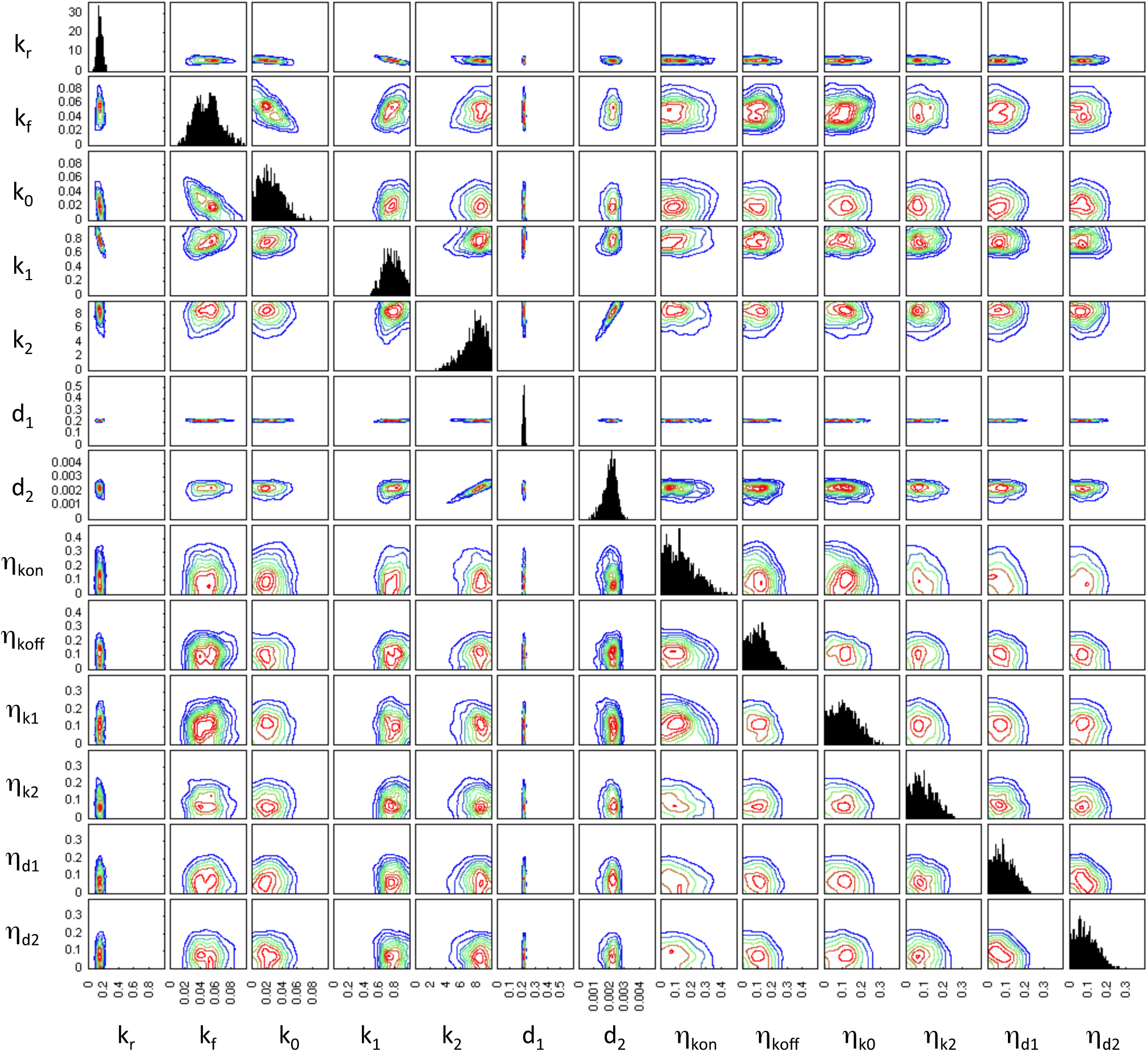
Relationships between the heaviest particles. Scatter plots of the particles with the highest weight in the final ABC-SMC population, shown for all pairs of parameters. Each point corresponds to a particle from the inferred posterior of one gene. Warmer hues are used to indicate a higher density of data points.

A comparison of Figues 3 and 4 suggests that a certain level of extrinsic noise is expected for all genes. However, the extrinsic noise may affect various combinations of rate parameters and it may not be possible to discern if, for example, the production rate or the degradation rate is more affected by extrinsic variability. While our inference procedure does not indicate a distinctive lower boundary for the amount of extrinsic noise affecting each reaction rate, there is usually an upper limit to the inferred noise parameters ranges. The extrinsic noise parameters for most genes are below 0.2 in the units set here (Figure 4); however, for some genes, *η_k_*_on_ and *η_k_*_off_ have relatively broad posterior marginal distributions.

To better understand the relationship between model parameters and observed patterns of gene expression, we look for correlations between means and variances of the inferred marginal parameters of each gene and the summary statistics used in the inference procedure (Figure 5). As expected, the correlation between the measured mRNA degradation rate, calculated form mRNA lifetime, and the inferred mRNA degradation rate parameter of the model, is close to one.

**Figure 5.**
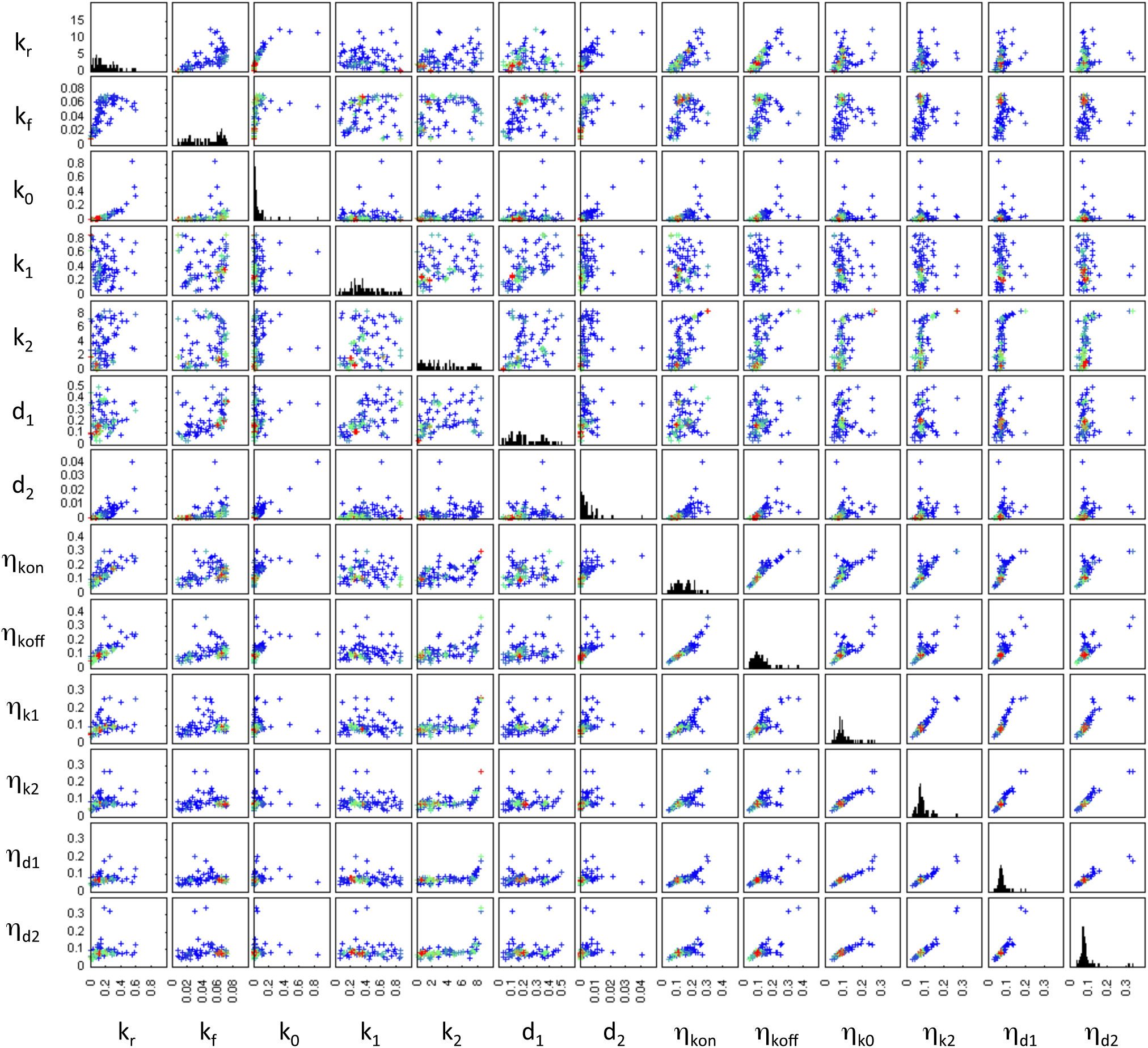
Heat maps of correlation coefficients between parameters and summary statistics. Heat maps are of the correlation coefficients calculated between experimentally obtained summary statistics and the mean (top) or the variance (bottom) of the marginal posterior for each model parameter. Correlation coefficients for which the associated p-values are greater than 0.05, after correcting for multiple testing using the Benjamini-Hochberg method,^39^ are treated as zero for plotting purposes.

The promoter switching rate parameters, *k*_on_ and *k*_off_, display positive and negative correlation with the mean mRNA number, respectively (as may be expected). They have the opposite relationship with the Fano factor associated with the mRNA distribution. This is consistent with the idea that distinct levels of transcription are required to account for the observed mRNA Fano factors. The corresponding extrinsic noise parameters *η_k_*_on_ and *η_k_*_off_ are positively correlated with mRNA abundance. However, the means and variances of the marginal distributions of these parameters are negatively correlated with the Fano factor of the mRNA distribution. This indicates that when promoter switching is affected by higher extrinsic noise, the mRNA distribution becomes more Poissonian as the effect of the two distinct promoter states is averaged out.

Curiously, the mean and variance of the protein degradation rate (*d*_2_) are positively correlated with mean mRNA number and negatively correlated with the mRNA Fano factor. Unlike the translation rate (*k*_2_), it shows no significant correlation with the mean or variance of the protein number.

### Parameter robustness and sensitivity

There are two complementary approaches to investigating the sensitivity of a modelled system to its parameters or inputs.^17^ One approach is to consider a single point in parameter space and study how the model responds to infinitesimal changes in parameters. This local approach usually involves calculating the partial derivatives of the model output with respect to the parameters of interest. Alternatively, one may consider how the model behaviour varies within a region of parameter space by sampling parameters and observing model behaviour. Regardless of the method used, different linear combinations of parameters will affect the model output to varying degrees.^18^ Gutenkunst *et al*^19^ coined the terms “stiff” and “sloppy” to describe these differences. They defined a Hessian matrix,

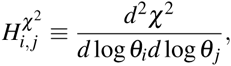

where *χ*^2^ provides a measure of model behaviour, such as the average squared change in the species time course. By considering the eigenvalues of this Hessian, *λ_i_,* the authors were able to quantify the (local) responsiveness of the system to a given change in parameters. Conceptually, moving along a stiff direction in parameter space causes a large change in model behaviour; conversely moving along a sloppy direction results in comparatively little effect on the output of the system.

Secrier *et al*^20^ later demonstrated how these ideas can be applied to the analysis of posterior distributions obtained by ABC methods.^21^ Principal component analysis (PCA) may be used to approximate the log posterior density using a multivariate normal (MVN) distribution. They showed that the eigenvalues of the covariance matrix, *s_i_,* of this MVN distribution are related to the eigenvalues of the Hessian as *λ_i_ =* 1*/s_i_.*

To assess the the stiffness/sloppiness of the inferred parameters we carry out PCA of the covariance matrices of log posterior distributions for each gene. In interpreting the results of the PCA we assume that the posterior distribution is, in practice, unimodal. The principal components (eigenvectors), *v*, and the corresponding loadings (eigenvalues), *s*, provided by the PCA are then used to obtain the eigen-parameters, *q*, as

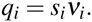

We calculate the projections of each parameter, *θ_i_*, onto each eigen-parameter, *q_j_,* as

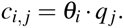

As a measure of the overall sloppiness of each parameter, *l*, we use the sum of the contributions of each parameter to the eigen-parameters, *l_i_* = Σ*_j_c_i_,_j_*. This can also be thought of as the sum of the projections of each principal component onto the parameter, weighted by the fraction of total variance explained by each of the principal components.

Having obtained a measure of the sloppiness of each parameter, for each gene, we carry out hierarchical clustering^22^ of genes and parameters using a Euclidean distance metric for both (Figure 6).

**Figure 6.**
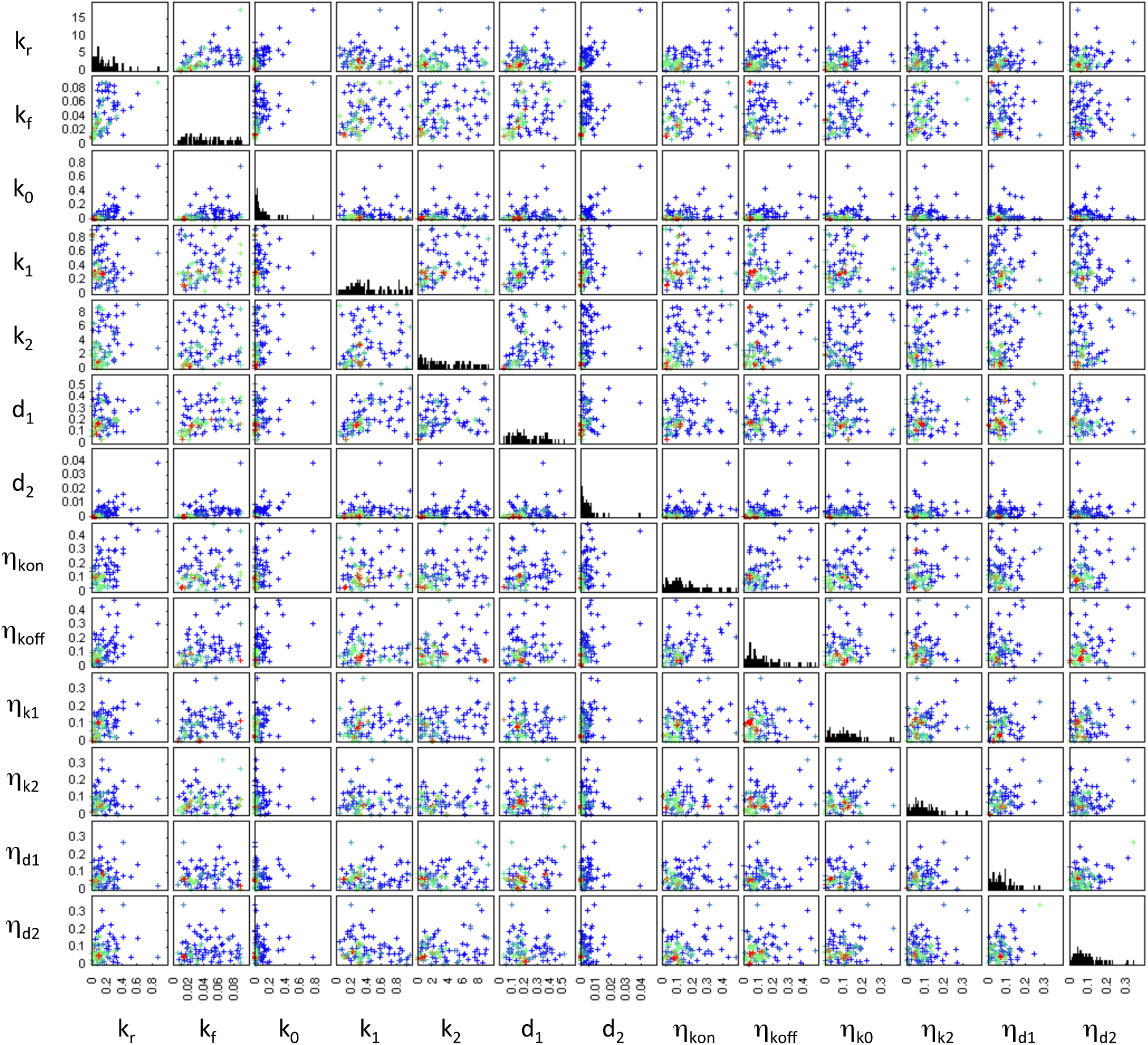
Clustering of genes and inferred posteriors according to parameter sloppiness. Clustergram showing a heat map of parameter sloppiness for each gene. Dendograms above and to the left of the heat map display the hierarchical tree obtained when clustering using a Euclidean distance metric.

The majority of genes show a similar pattern of parameter stiffness/sloppiness. The most distinctive and the second most distinctive clusters consist of just two genes each, *yiiU* with *aceE* and *cspE* with *map,* respectively. These four genes are distinguished by unusually sloppy promoter activity ratio, *k_r_*, and promoter switching frequency, *k_f_*, parameters. The pair *yiiU* and *aceE* display a high ratio of protein variance to protein mean (Fano factor) and are stiff with regard to the protein degradation rate noise parameter 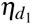. *cspE* also has a high Fano factor of the protein distribution while *map* has an unusually low mRNA Fano factor. What these four genes appear to have in common is that the variability in their protein numbers is difficult to explain based solely on the mRNA variability. Thus, a higher level of extrinsic noise is inferred to account for the observed variability. Since these genes comprise a small minority, it may be that their expression is subject to regulatory mechanisms that are not well approximated by the two state model. The remaining majority of genes are broadly divided into two similar groups which differ mostly in the sloppiness of *k*_0_.

The noise and rate parameters segregate into two clusters with the noise parameters generally being sloppier than the rate parameters (Figure 6). The least sloppy parameter is the mRNA degradation rate (*d*_1_). This is not surprising since it was used, together with the molecule number summary statistics, to infer the posterior distribution. Of the rate parameters, the basal transcription rate (*k*_0_) is the sloppiest and often approaches the noise parameters in its sloppiness. Since this parameter is defined as a fraction of the active transcription rate (*k*_1_), its relative sloppiness should not be equated to a lack of importance. For most genes the marginal posterior of *k*_0_ is largely constrained to the lower half of its prior distribution, *U*(0,1). The only exception being the gene *map* for which the measured mRNA Fano factor was close to one and the marginal posterior of *k*_0_ is in the top half of the prior range. The mean of the marginal posterior of *k*_0_ is negatively correlated with the mRNA Fano factor across all genes (Figure 5). The two other parameters that influence the mRNA Fano factor, *k_r_* and *k_f_*, are the next sloppiest rate parameters.

## Conclusions

Cell-to-cell variability in genetically homogeneous populations of cells is a ubiquitous phenomenon.^23–25^ Attempts to quantify it are complicated by the difficulty of assigning it to a single cellular process or any one experimentally measurable variable. It can also be difficult, for example, to distinguish between the intrinsic stochasticity of biochemical processes in the short term and longer term variations which may have been inherited from previous cell generations.

By including a representation of extrinsic noise in our model of gene expression we infer the extent to which the rates of biochemical processes can vary between cells while still producing the experimentally measured mRNA and protein variability. We demonstrate the usefulness of an efficient method for exact stochastic simulation of the two-state model of gene expression. This model is necessary to explain the experimentally measured mRNA variation (Fano factor) and is capable of describing the majority of the observed data. We show that the amount of extrinsic noise affecting most genes appears to be limited, but non-negligible.

The exact simulation method described here occupies a niche between those cases when only samples from the steady state mRNA distribution of the two-state model^3,26,27^ are required, and cases when an approximation to the protein distribution^13,28^ is sufficient. The computational advantages of the simulation method described here are limited to specific conditions, such as, low numbers of mRNA molecules and higher numbers of protein molecules. The most limiting factor of this simulation method is that it is not applicable to models in which the protein products affect upstream processes such as promoter activity, transcription or translation. The addition of such interactions would mean that the assumptions used in deriving the Poissonian relationship between the number of surviving protein molecules produced form a given mRNA molecule and mRNA’s lifetime would no longer be satisfied. Perhaps an approximate algorithm could be developed on the basis of algorithm (1) to handle such situations. Alternatively, the tau-leaping algorithm,^29^ or moment expansion,^30,31^ may be more appropriate for models involving these kinds of feedback interactions. Algorithm (1) could, however, be naturally extended to models involving regulatory interactions between non-coding RNAs as the simulation of that part of the model is equivalent to Gillespie’s exact algorithm. Although here we use summary statistics of mRNA and protein number measurements, the simulation method is also applicable to cases where a direct comparison between sample distributions, for example using the Hellinger distance, is required.

The inferred extrinsic noise parameters will also include the effects of regulatory mechanisms that are not well described by the two-state model. In this sense, our definition of noise becomes blurred with our ignorance about the regulatory interactions involved in the expression of each gene. Nonetheless, the biochemical mechanisms governing gene expression in a given species are shared between many genes. This is in agreement with our observation that, for most genes, inferred model parameters show similar patterns of sloppiness. If we are able to refine our understanding of the shared aspects of gene expression, we may be able to improve our understanding of both the nature of the noise affecting it, and the regulatory mechanisms controlling it. In practice this may mean finding a mechanistic explanation for the two-state model or further refining it to achieve a better agreement between simulations and experimental results.

The *in silico* approach used here not only relied on, but was inspired by the experimental work of Tanaguchi *et al*.^7^ As the resolution of high throughput experimental techniques and the quantity of data they generate continues to increase, more complete observations of cellular processes may begin to yield data amenable to statistical analysis and inference of extrinsic noise. These may in turn require other modelling, computational and theoretical approaches which would not rely on the assumptions and simplifications that we make in this work.^32^

## Methods

### 1 Modelling gene expression

A simple model of gene expression may represent the processes of transcription and translation using mass-action kinetics to describe production and degradation of various species as pseudo-first order reactions. Such a model may be simulated stochastically to take into account the intrinsic variability of processes involving low numbers of molecules. In the simplest version of this model, mRNA is produced from the promoter at a constant rate. However, such Poissonian mRNA production is often not sufficient to account for the variability in mRNA numbers measured experimentally in both prokaryotic and eukaryotic cells. In addition to this, for many genes, transcription appears to occur in bursts rather than at a constant rate. These characteristics of gene expression have been observed in organisms as diverse as bacteria,^7^ yeast,^4^ amoeba^2^ and mammals.^3^ One model of gene expression that takes this into account is the, so called, two-state model.

#### 1.1 The two-state promoter model

In the two-state model of gene expression, a gene’s promoter is represented as either active or inactive.^5,16^ Here we use a variant of the two-state model with the inactive state corresponding to a lower transcription rate rather than no transcription at all. For each state of the promoter, transcription events at that promoter are represented by a Poisson process with rate parameter corresponding to the transcription rate. Biochemical processes such as transcription factor binding or reorganisation of chromatin structure may account for the existence of several distinct levels of promoter activity. However, which factors play a dominant role in the apparent switching, remains an unanswered question.

The Gillespie algorithm^33^ may be used to simulate all the reactions represented by this model and obtain a complete trajectory of the system through time. However, in this case we are only interested in the number of molecules present at the time of measurement. We use a model-specific stochastic algorithm (Algorithm 1) which allows us to reduce the number of computational steps required to obtain a single realisation from the model.

The following reactions, represented using mass-action kinetics, comprise the two-state model:

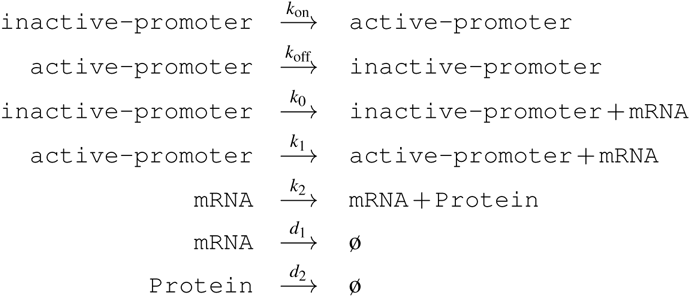

The propensity functions (hazards) for each of the above reactions are listed below:

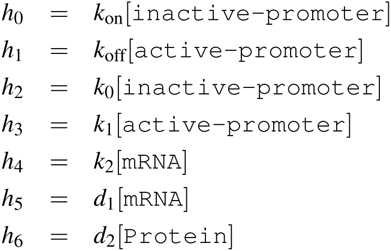

Here the square brackets refer to the number of molecules of a species rather than its concentration.

The model presented here relies on a number of assumptions about the process of gene expression. Firstly, that the production of mRNA and protein can be described sufficiently well by pseudo-first-order reactions. Secondly, that degradation of mRNA and protein can be described as an exponential decay. In a bacterial cell, mRNA molecules are degraded enzymatically and typically have a half-life on the scale of several minutes. The half-life of protein molecules usually exceeds the time required for cell growth and division during the exponential growth phase. Thus, dilution due to partitioning of protein molecules between daughter cells tends to be the dominant factor in decreasing the number of protein molecules. Here we do not build an explicit model of cell division, instead the decrease in protein numbers is approximated by an exponential decay. Finally, it is assumed that there is no feedback mechanism by which the number of mRNA or protein molecules produced by the gene affects its promoter switching, transcription or translation rates.

##### 1.1.1 Representing extrinsic noise

We model extrinsic noise by perturbing the reaction rate parameters, using a Gaussian kernel, before each simulation of the model.^34^ The effect of extrinsic noise on each reaction is assumed to be independent. The reaction rates associated with a particular gene are termed nominal parameters (*θ_n_*).

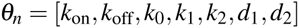

The values determining the magnitude of the perturbation are termed the noise parameters (*η*).

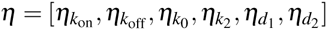

Together they comprise the full parameter set for the model *θ* = [*θ_n_*, *η*].

In the case of the two-state model of a single gene, each *θ_n_* has a corresponding extrinsic noise parameter with the exception that the basal transcription rate 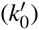 is defined as a fraction of the active transcription rate 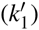 so the two reaction rates are subject to the same perturbation 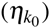 before each simulation. This is motivated by the idea that extrinsic factors affecting the transcription rate do not depend on the state of the promoter. The parameters used to generate a single realisation from the two-state model are obtained by sampling from *f*(*μ*, *σ*). Where *f* is a truncated normal distribution, restricted to non-negative values by rejection sampling, with *μ* and *σ* being the mean and standard deviation of the corresponding normal distribution.

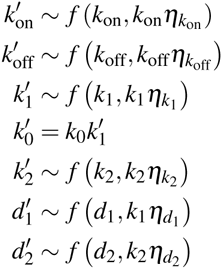

The final time point of each simulation represents the number of mRNA and protein molecules in a single cell at the time of measurement.

#### 1.2 Simulation procedure

In order to reduce the computational cost of each simulation, rather than using Gillespie’s direct method to simulate the entire trajectory of mRNA and protein numbers, we employed Algorithm 1 to obtain samples of the numbers of mRNA and protein molecules at the time of measurement (*t_m_*). First, a realisation of the telegraph process is used to obtain the birth and decay times of mRNA molecules. These are then used to sample the number of protein molecules that were produced from each mRNA molecule and survived until *t_m_*. This procedure makes use of the Poisson relationship between the life time of an individual mRNA molecule and the number of surviving protein molecules that were produced from it. This relationship is derived in Appendix 2.2 and its use is illustrated in Figure 7. The final result is the number of both mRNA (*M*) and protein (*P*) molecules present in the system at *t_m_*.

**Figure 7.**
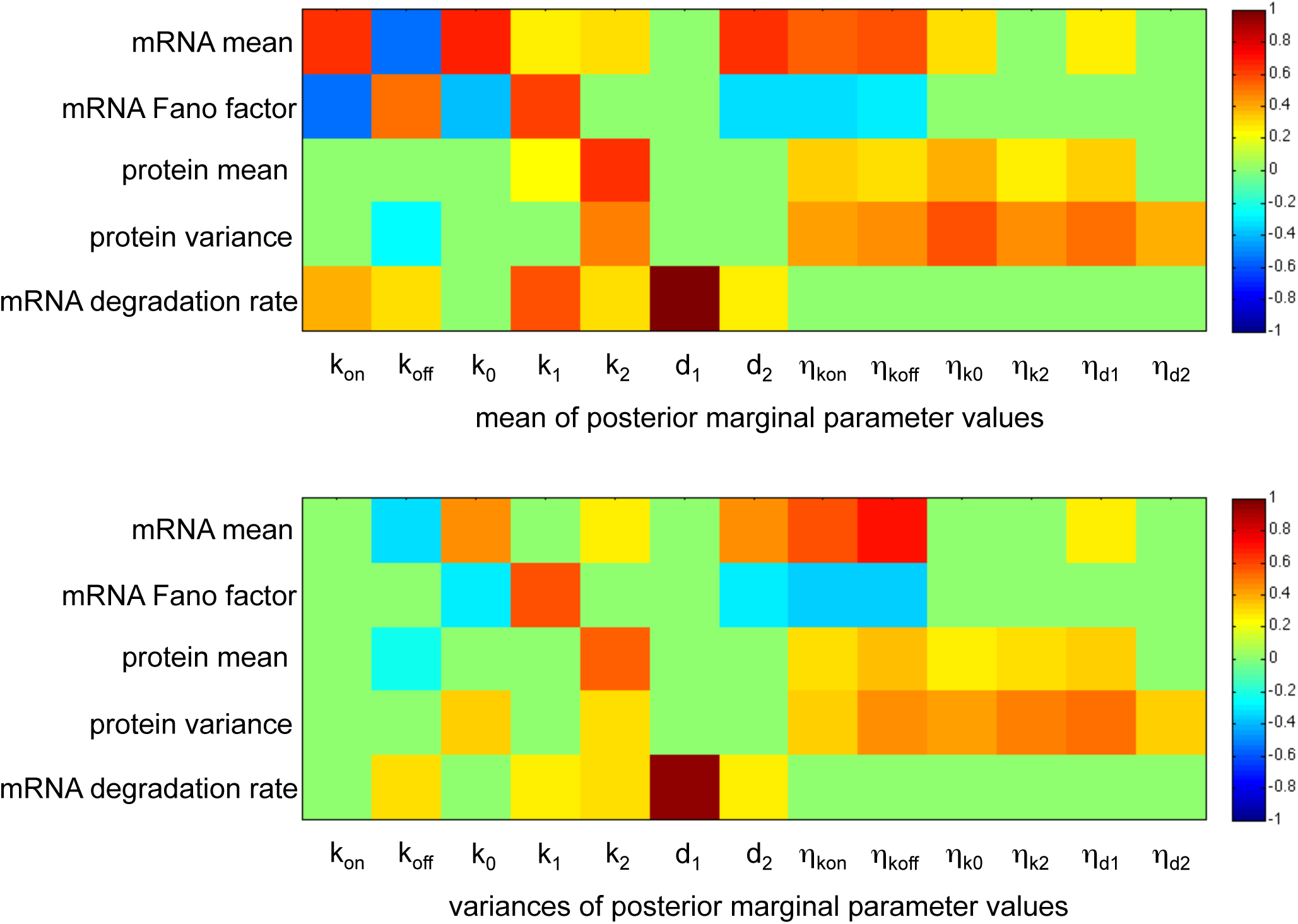
Illustration of the principle behind Algorithm 1. An illustration of how the birth and death times of an mRNA molecule are used to obtain the number of proteins that were produced from it and then survived until the time at which mRNA and protein numbers were measured.

#### 1.3 Use of experimental data

Using an automated fluorescent imaging assay, Taniguchi et al^7^ were able to quantify the abundances of 1018 proteins from a yellow fluorescent protein fusion library. We focus on a subset of 87 genes from the published data set from.^7^ These are all the genes for which, in addition to protein numbers, the experimental data include both fluorescence *in situ* hybridization measurements^35^ of mRNA numbers and mRNA lifetimes measurements obtained using RNAseq.^36^ We note that these genes are not a random sample from the set of all genes and exhibit higher than average expression levels.

To identify model parameters for which the two-state model, with extrinsic noise, is able to reproduce the experimental measurements, we carry out Bayesian inference using an ABC sequential Monte Carlo (SMC) algorithm that compares summary statistics from simulated and experimental data.^37^ Specifically we used the following summary statistics: (1) the mean numbers of mRNA molecules; (2) the Fano factors of mRNA molecule distributions; (3) the mean numbers of protein molecules; (4) the variances of protein molecule numbers; and (5) mRNA lifetimes converted to expontial decay rate parameters. The distributions of these summary statistics are shown in Figure 8. We assume that the summary statistics correspond to steady state expression levels for each gene. While there is no guarantee that this is the case for every gene, the majority of genes are unlikely to be undergoing major changes in their expression level given that the cells are in a relatively constant environment.

**Figure 8.**
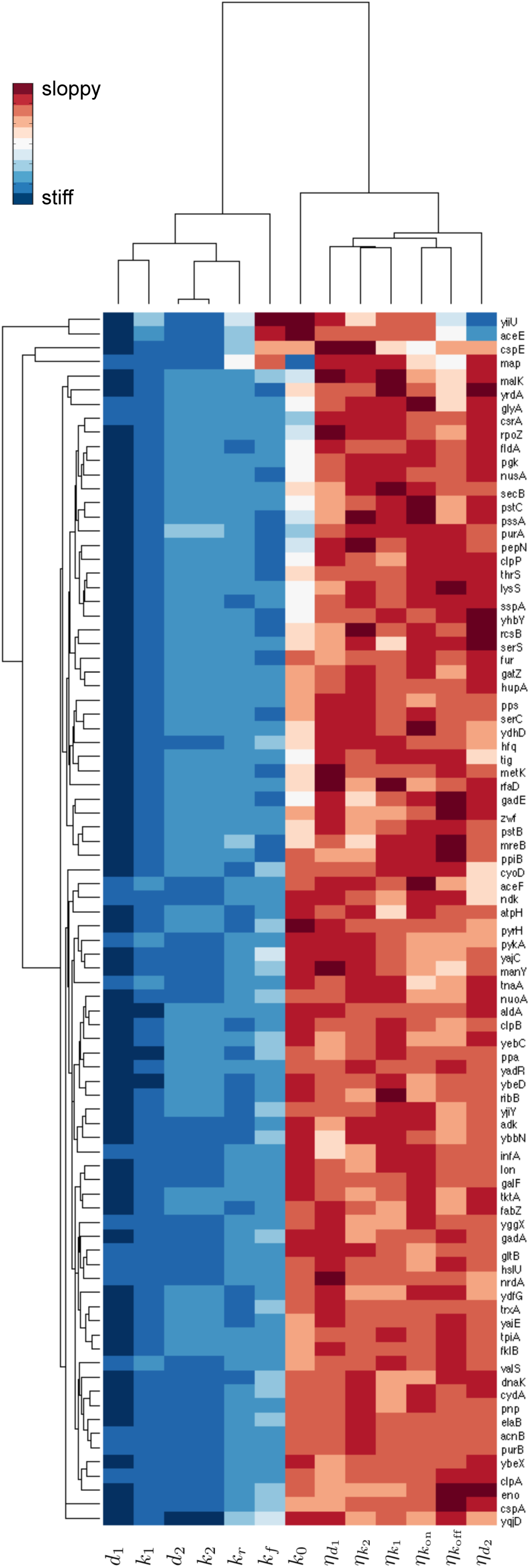
Experimentally measured summary statistics. Each point on the scatter plots is an estimate of the corresponding summary statistic or mRNA lifetime from experimental measurements.

Taniguchi *et al*^7^ used images of about a thousand cells to obtain estimates of mean mRNA numbers, mRNA Fano factors, mean protein numbers and protein number variances. For this reason, we use 10^3^ simulation runs when calculating summary statistics. The experimental measurements of mRNA lifetimes are compared directly to the mRNA degradation rate parameter (*d*_1_) in the model by assuming that lifetimes correspond to the inverse of the decay rate.

#### 1.4 Inference procedure

We use an ABC-SMC algorithm to infer plausible parameter sets for the two-state model based on the experimental data. The inference procedure is similar to that employed by,^21,37,38^ as described in Algorithm 2.

For the distance metric, *d*, we take the Euclidean distance between the logarithms of each type of experimental measurement (*D_i_*) and the corresponding simulation results (*x_i_*):

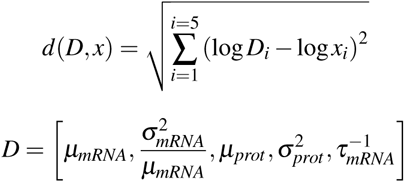

Where *μ_mRNA_* is the mean number of mRNA molecules; 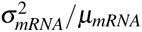 is the Fano factor of the mRNA distribution; *μ_prot_* is the mean number of protein molecules; 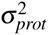 is the variance of the protein distribution and 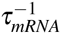 gives the exponential decay rate constant for mRNA degradation based on the measured mRNA lifetime (τ_mRNA_).

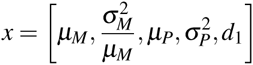

Where *μ_M_* is the mean number of mRNA molecules; 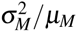 is the Fano factor of the mRNA distribution; *μ_P_* is the mean number of protein molecules; 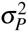 is the variance of the protein distribution and *d_1_* corresponds to the nominal mRNA degradation rate. The first sampled population of particles (population zero in Algorithm 2), provides a benchmark for the choice of *ε* values in the next population. Since we have no knowledge of the distribution of distances until a set of particles is sampled, all particles are accepted in the first population. For subsequent populations, ε values are chosen such that the probability of acceptance with the new *ε* value is equal to *q_t_*. The vector *q* is chosen prior to the simulation. This allows for larger decreases in *ε* in the first few populations while keeping the actual epsilon values used, a function of the distances (*g*) in the previous population. New populations are sampled until the final epsilon value is reached ε*_f_* = 0.1. To obtain *θ*\* from *θ* we use a uniform perturbation kernel:

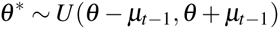

where *μ_t−1_* is the vector of standard deviations of each parameter in the previous population.

##### Algorithm 1 Simulation of the two-state model

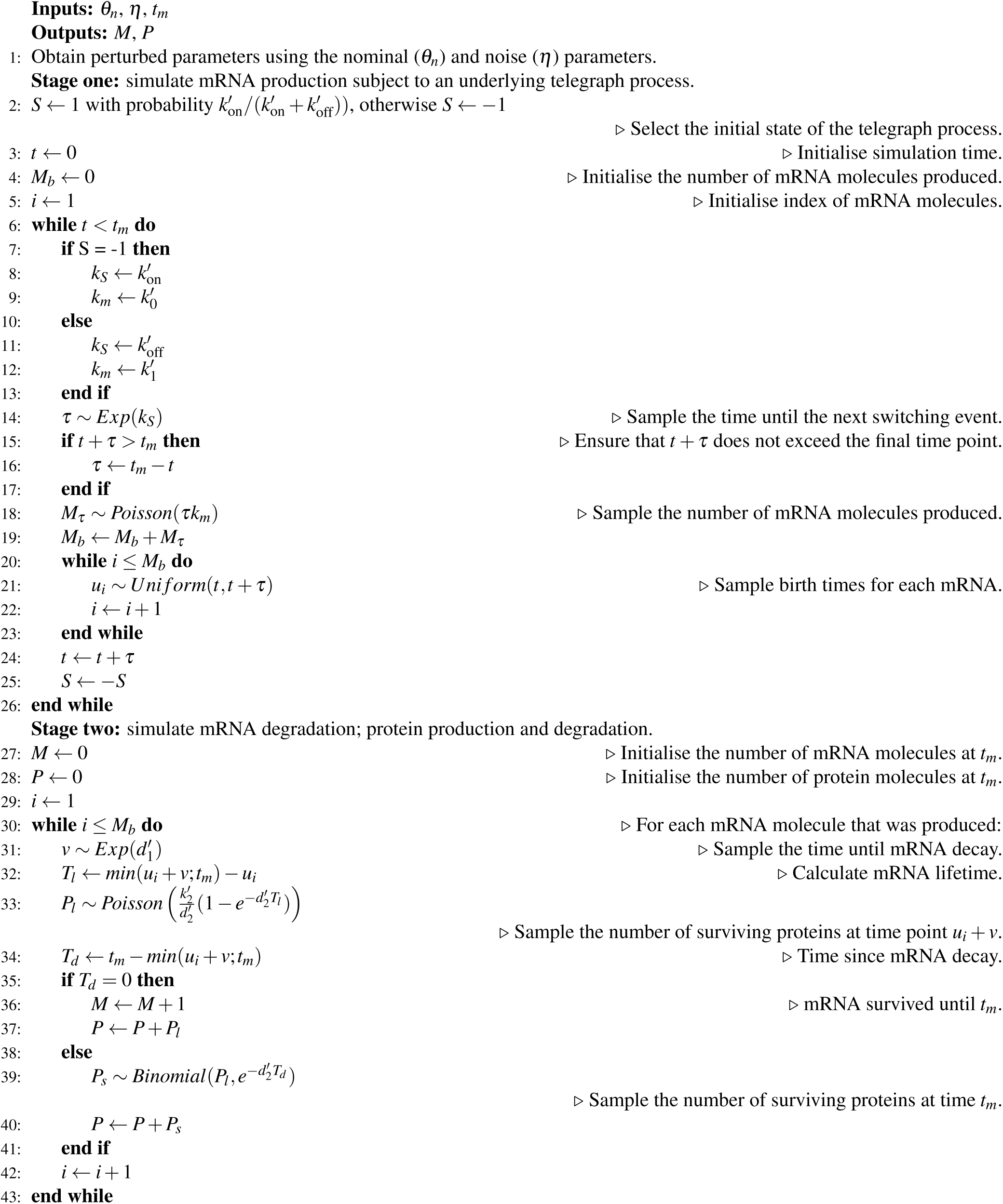

##### Algorithm 2 ABC-SMC with summary statistics

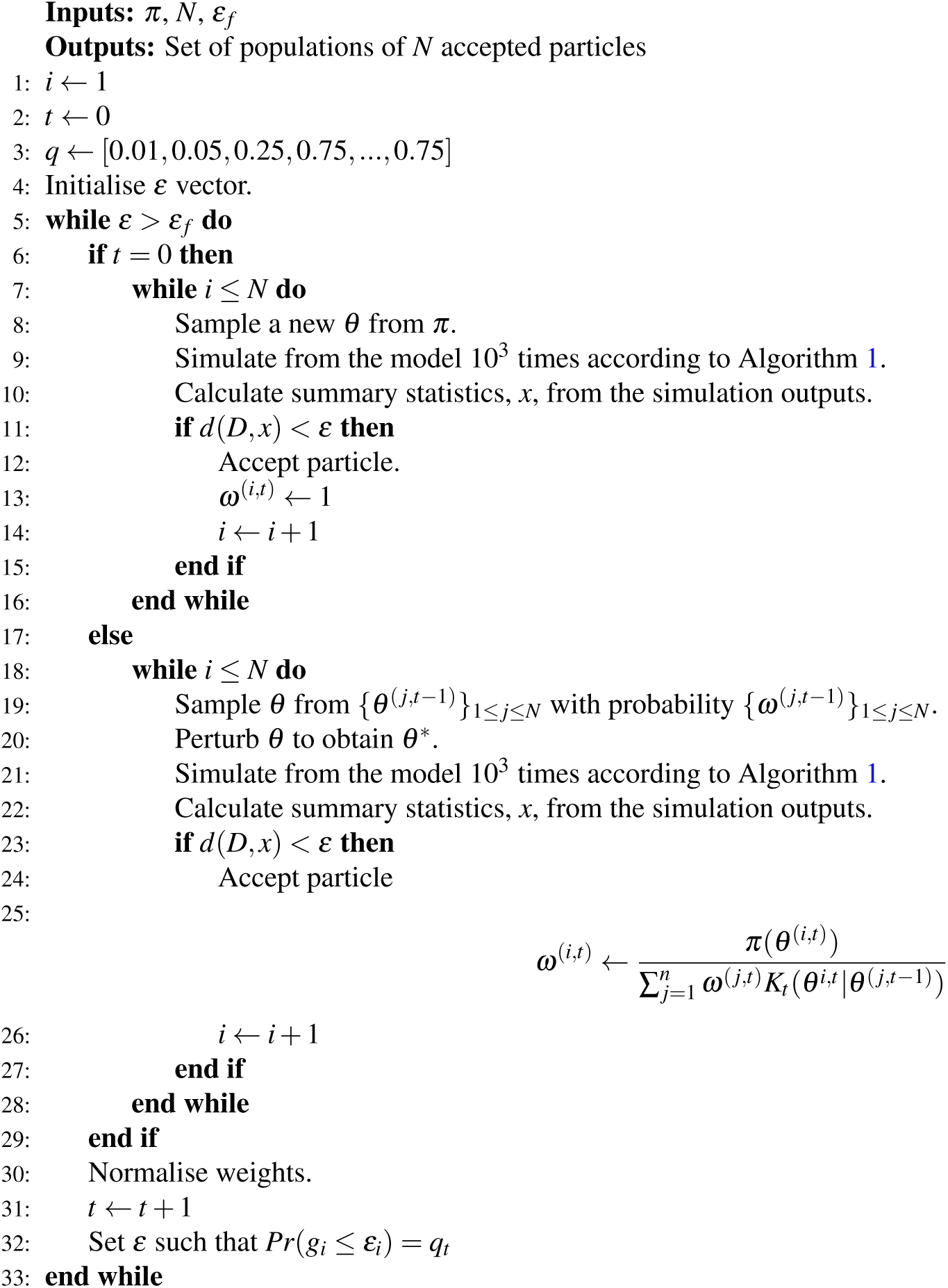

##### 1.4.1 Parameter prior

The telegraph process may be parametrized in terms of the ratio of probabilities of switching events (*k_r_*) and the overall frequency with which events occur (*k_f_*):

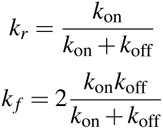

To obtain *θ*, the vector of parameters used in the ABC-SMC inference procedure (Algorithm 2), rate and noise parameters are sampled from the following uniform priors,

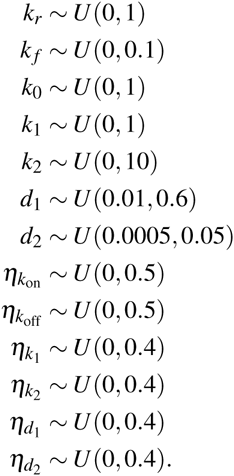

The parameters for the telegraph process, sampled from the prior as *k_r_* and *k_f_*, are converted to *k*_on_ and *k*_off_ before being passed to the simulation algorithm (Algorithm 1) as follows,

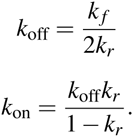

Rate parameters *k_r_* and *k*_0_ as well as the noise parameters (*η*) are unit-less. The remaining parameters have units 1*s*^−1^.

To ensure that *M* and *P* are from a distribution close to equilibrium, simulation duration is set depending on the nominal degradation rates for mRNA (*d*_1_) and protein (*d*_2_),

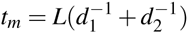

where *t_m_* is the final time point and *L* is a constant chosen arbitrarily to indicate the desired proximity to the steady state distribution. Here we use *L* = 5.

## Author’s contributions

O.L., P.K. and M.P.H.S. designed the study. O.L. carried out the computational work. O.L. and M.P.H.S. wrote the paper. All authors read and reviewed the final paper.

## Additional Files

### Appendix — Derivation of the Poissonian relationship between the number of surviving protein molecules and mRNA lifetime

#### 2 Derivations

Consider the following system of reactions.

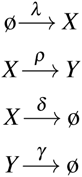

The system is allowed to run, starting with zero tokens of *X* and *Y*, from time *t*_0_ until time *t_m_*. The following table lists events of interest which can occur in the system and the symbols used to represent the number of times each even occurs in the time interval *T = t_m_* −*t*_0_.

| Symbol | Event being counted |
| --- | --- |
| $i$ | $X$ is produced |
| $j$ | $X$ survived until time $t_m$ |
| $\hat{i}$ | $X$ does not survive until time $t_m$ (i.e. $X$ is degraded or converted to $Y$) |
| $\hat{j}$ | $X$ is degraded |
| $k$ | $X$ is converted into $Y$ |
| $l$ | $Y$ survives until time $t_m$ |

##### 2.1 Deriving probability distributions for the numbers of events

A standard result is that the number of events which occur over a time period will have a Poisson distribution if the events are occurring with a uniform probability in a time interval. This applies to the production of *X*.

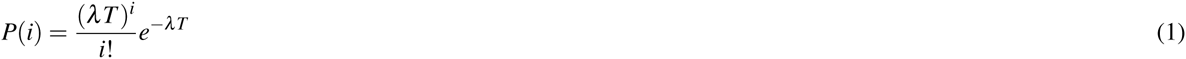

Where *P*(*i*) is the probability that *X* is produced *i* times during the time period *T*.

##### 2.2 Finding the number of surviving tokens of *X*

In order to find the probability distribution of the number of *X* tokens after the time period *T* we consider the probability that *j* tokens of *X* remain given that *i* tokens of *X* were produced.

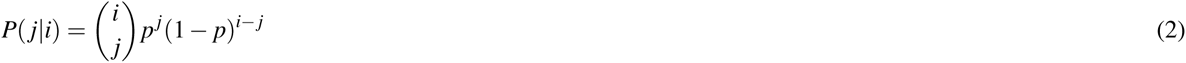

Where *p* is the probability that *X* survives given that it was produced during the time period *T*.

Let *P*(*j*) be the probability that *j* tokens of *X* were produced during *T* and survived regardless of the number (*i*) of *X* tokens produced.

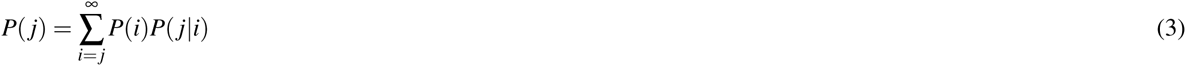

Using equation 1 and 2.

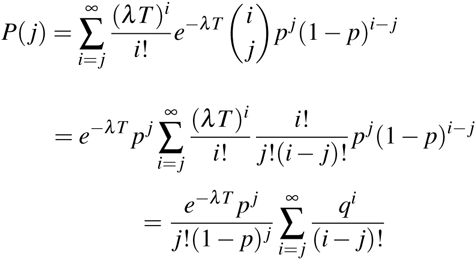

Where *q = λT*(1 − *p*).

Considering just the summation term.

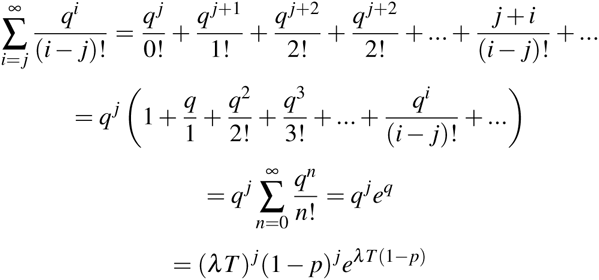

Thus, the expression for *P*(*i*) can be written as:

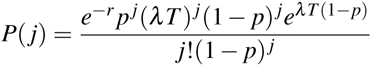

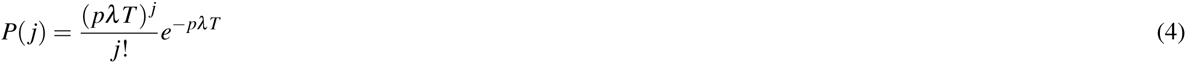

*P*(*j*) has a Poisson distribution with the parameter *pλT*.

Next, we need to find *p*. The probability that *X* was produced during a infinitesimally short time interval *dt* is given by *dt*/*T*. That is to say that *X* is produced once at some point during *T* and the probability of *X* production is uniform over *T*. The probability of a token of *X* produced at time *t* surviving until time *t_m_* follows an exponential decay with a rate *δ*′. In this case *δ*′ = *ρ* + *δ*. Thus the probability of a token of *X* being produced at time *t* and surviving until *t_m_* is given by

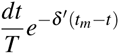

The overall probability that *X* survives given that it was produced can be found by integrating over the time period *T*.

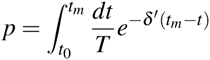

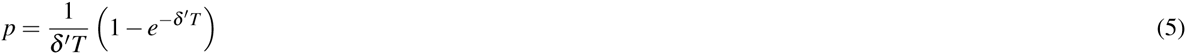

Where *T = t_m_ −t*_0_.

The numbers of the other events are also Poisson distributed. Similarly to *P*(*j*), the number of tokens of *X* which did not survive 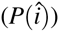 will also be Poisson distributed.

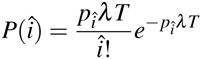

Where 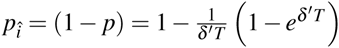.

Also, for *P*(*k*), the distribution of the number of tokens of *Y* produced

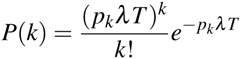

with 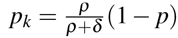.

